# IFITM3 restricts human metapneumovirus infection

**DOI:** 10.1101/290957

**Authors:** Temet M. McMichael, Yu Zhang, Adam D. Kenney, Lizhi Zhang, Ashley Zani, Mijia Lu, Mahesh Chemudupati, Jianrong Li, Jacob S. Yount

## Abstract

Human metapneumovirus (hMPV) utilizes a bifurcated cellular entry strategy, fusing either with the plasma membrane or, after endocytosis, with the endosome membrane. Whether cellular factors restrict or enhance either entry pathway is largely unknown. We found that the interferon-induced transmembrane protein 3 (IFITM3) inhibits hMPV infection to an extent similar to endocytosis-inhibiting drugs, and an IFITM3 variant that accumulates at the plasma membrane in addition to its endosome localization provided increased virus restriction. Mechanistically, IFITM3 blocks hMPV F protein-mediated membrane fusion, and inhibition of infection was reversed by the membrane destabilizing drug amphotericin B. Conversely, we found that infection by some hMPV strains is enhanced by the endosomal protein Toll-like receptor 7 (TLR7), and that IFITM3 retains the ability to restrict hMPV infection even in cells expressing TLR7. Overall, our results identify IFITM3 as an endosomal restriction factor that limits hMPV infection of cells.

## INTRODUCTION

Human metapneumovirus (hMPV) is a member of the *Pneumoviridae* family, a recent classification distinguishing it from related *Paramyxoviridae* family members[1]. Since its discovery in 2001[2], hMPV is increasingly recognized as a significant cause of respiratory infections in infants, children, and the elderly[3–6]. This virus infects over 85% of the population by age ten[7], and is one of the most common causes of respiratory tract infections in infants[3, 5]. The frequency of detection of hMPV in elderly patients with respiratory infections approaches that of influenza virus[4]. However, unlike influenza virus, there are currently no licensed vaccines or drugs targeting hMPV. While vaccination strategies have been proposed for eliciting an adaptive immune response against hMPV[8, 9], roles for innate immune restriction factors in controlling the virus remain to be identified.

The early classification of hMPV as a paramyxovirus suggested that it would exhibit pH-independent fusion at the plasma membrane similarly to other prototypical paramyxoviruses, such as Sendai virus (SeV)[10]. However, early reports indicated that acidic pH enhanced fusion mediated by the F proteins of certain hMPV strains[11, 12]. Follow up studies utilizing single virus particle tracking and fusion assays showed that a portion of hMPV fuses at the plasma membrane as is typical of paramyxoviruses, but also that a significant fraction of the virus is endocytosed and fuses with endosomal membranes in a manner characteristic of low pH-dependent viruses, such as influenza A virus (IAV)[13–15]. These data suggested that hMPV utilizes a bifurcated cellular entry strategy with characteristics typified by both SeV and IAV. From this, we reasoned that hMPV may be susceptible to inhibition by interferon-induced transmembrane protein 3 (IFITM3), which is a cellular antiviral factor that inhibits virus-endosome membrane fusion reactions[16–21].

IFITM3 broadly restricts endocytosed virus infections by blocking their fusion, thus preventing cytosolic entry of virus genomes[16]. Though the exact mechanism of fusion inhibition is unknown, IFITM3 may alter membrane properties, including the rigidity and curvature of endosome membranes[18, 22]. Our recent work demonstrated that a palmitoylated amphipathic helix within IFITM3 is required for its inhibition of virus protein-mediated membrane fusion[23–25]. Viruses such as IAV that enter cells primarily through endocytosis are strongly inhibited by IFITM3 while viruses such as SeV that fuse at the cell surface are poorly inhibited[26, 27]. IFITM3 KO mice and humans with deleterious IFITM3 gene polymorphisms experience severe influenza virus infections, confirming the importance of this protein in antiviral defense *in* vivo[28–30]. Effects of IFITM3 on hMPV have not yet been adequately investigated.

Here we show that IFITM3 is able to inhibit hMPV infection, thus identifying the first known cellular restriction factor for this important respiratory pathogen. We observed partial inhibition of hMPV infection by endocytosis inhibitors and partial restriction by IFITM3, both of which are consistent with the dual entry mechanism proposed for this virus. We demonstrate that IFITM3 can block membrane fusion mediated by the hMPV F protein and, importantly, that manipulating the levels, localization, or activity of IFITM3 in cells can have significant effects on hMPV infection.

## METHODS

### Cell culture, transfections, and transductions

HEK293T, A549, Vero, and MEF cells were grown in DMEM supplemented with 10% Equafetal FBS (Atlas biologicals). LLC-MK2 cells were grown in OPTI-MEM reduced serum media + Glutamax supplement (Thermo Fisher) with 2% FBS (Atlas biologicals). HAP1 WT and IFITM3 KO cells (Horizon Discovery) were grown in IMDM with 10% Equafetal FBS. Macrophages were grown in RPMI with 10% Equafetal FBS. All cells were grown at 37°C with 5% CO_2_ in a humidified incubator. HEK293T cells transduced with human TLR7 or vector control were purchased from InvivoGen. NEDD4 WT and KO MEFs were generated by Dr. Hiroshi Kawabe (Max Planck Institute)[31, 32]. Macrophage cell lines were generated by Dr. Douglas Golenbock (University of Massachusetts) and obtained through the NIH-sponsored BEI Resources. Cells were transfected with plasmids using LipoJet transfection reagent (Signagen Laboratories) following the manufacturer’s instructions. For generation of stable cell lines, myc-IFITMs were inserted into pLenti-puro, and VSV G pseudotyped lentiviruses were generated as described previously[33].

### Treatment with IFN, siRNA, or drugs

Where indicated, cells were treated with human IFN-β (obtained through BEI resources and utilized at a 1:100 concentration) or with IFN-α2 (eBioscience, 1:1000 concentration) for 24 h to induce IFITM3 expression. IFITM3 knockdown was achieved using Dharmacon ON-TARGET Plus Smart Pool human IFITM3-targeting (L-014116) or non-targeting control (D-001810-10-20) with Lipofectamine RNAiMAX reagent (Invitrogen) according to the manufacturer’s protocol. For experiments involving inhibitors or drugs, cells were treated with chlorpromazine or genistein at concentrations of 10μg/mL or 200μM, respectively, or with 2.5 ug/mL amphotericin B.

### Virus propagation, infection, and flow cytometry

Influenza A virus A/PR/8/34 (H1N1, PR8) was propagated in 10 day old embryonated chicken eggs (Charles River) for 48 h at 37C and titered on MDCK cells. SeV expressing GFP was generated by Dr. Dominique Garcin (University de Geneve), and was propagated in 10 day old embryonated chicken eggs at 37°C for 40 h and titered on Vero cells. hMPV and hMPV-GFP were generated by a reverse genetics system based on the NL/1/00 (A1) strain utilizing previously described methodology[34]. The hMPV 02-202 (B1) strain was provided by Dr. John Williams (University of Pittsburgh). All hMPV strains were propagated in Vero cells, concentrated by ultracentrifugation through a 20% sucrose cushion, and titered on LLC-MK2 cells. For flow cytometry quantification of infection, IAV-infected cells were stained with anti-H1N1 IAV NP (obtained from BEI resources), hMPV-infected cells were stained using anti-hMPV antibody (Millipore, MAB80138), and cells infected with GFP-expressing viruses were analyzed for GFP fluorescence directly. Flow cytometry was performed on a FACSCanto II flow cytometer (BD Biosciences), and analyzed using FlowJo software.

### Western blotting and antibodies

For Western blotting, cells were lysed in buffer containing 0.1 mM triethanolamine, 150 mM NaCl, and 1% SDS at pH 7.4 supplemented with EDTA-free Protease Inhibitor Cocktail (Roche) and Benzonase Nuclease (Sigma). Primary antibodies for IFITM1 (Cell Signaling), IFITM2 (Cell Signaling), IFITM3 (Proteintech group), HA tag (HA.11, Biolegend), GAPDH (Invitrogen), Actin (Abcam), Tubulin (Antibody Direct), Myc tag (Developmental Studies Hybridoma Bank at the University of Iowa, deposited by Dr. J. Michael Bishop, catalog no. 9E10), and NEDD4 (Millipore) were used at 1:1000 dilutions.

### Cell-cell fusion assay

The trypsin-independent hMPV F protein was cloned previously by mutating the RQSR motif of the hMPV NL/1/00 strain[35]. Cell-cell fusion assays were performed as outlined previously[24, 36]. For pH 5.0 pulses, freshly prepared DMEM containing 25 mM MES at pH 5.0 was used to replace cell media for 2 min. Cells were then washed with PBS and incubated in standard media for 8 hours at 37°C. Luciferase activity was measured using the Promega Dual-Luciferase Reporter Assay System.

## RESULTS

### Overexpression of IFITM3 inhibits hMPV infection

To examine effects of IFITM3 on hMPV strain NL/1/00 (A1) infection, we compared susceptibility of HEK293T cells after stable transduction with empty vector or myc-tagged human IFITM3 (Figure 1A). Importantly, HEK293T cells do not express detectable IFITM3 or other IFITMs at baseline and are thus a useful model for assessing virus infection upon introduction of IFITM3 (Supplementary Figure 1A)[37]. Percent hMPV infection of IFITM3-expressing cells as measured by flow cytometry was decreased by approximately half compared to vector control cells (Figure 1B). As a positive control, we tested susceptibility of the cells to IAV and saw that infection was reduced almost completely in IFITM3-expressing cells (Figure 1A)[16, 24]. As a negative control, we found that SeV was largely insensitive to expression of IFITM3 (Figure 1A)[26, 32]. These results are consistent with the endocytic entry of IAV, plasma membrane entry of SeV, and dual entry of hMPV. To confirm the role of endocytosis in hMPV infection of HEK293T cells, we examined effects of two broad inhibitors of endocytosis, chlorpromazine and genistein. These inhibitors caused a nearly complete elimination of infection of cells by IAV and blocked hMPV infection by roughly 50%, correlating in magnitude to the extent of inhibition by IFITM3 for both viruses (Figure 1B,C). We additionally observed that overexpression of IFITM1 partially inhibited hMPV infection of HEK293T cells, and that IFITM2 modestly inhibited infection, though both proteins strongly inhibited IAV infection (Supplementary Figure 1). These results suggested that IFITMs 1-3 are capable of inhibiting hMPV when overexpressed. However, given that IFITMs 1 and 2 do not compensate for loss of IFITM3 *in vivo*[28–30, 38, 39], and given that our early studies examining IFITM3 KO cells suggested an important role for IFITM3 in restricting hMPV, we focused our subsequent experiments on IFITM3.

**Figure 1:**
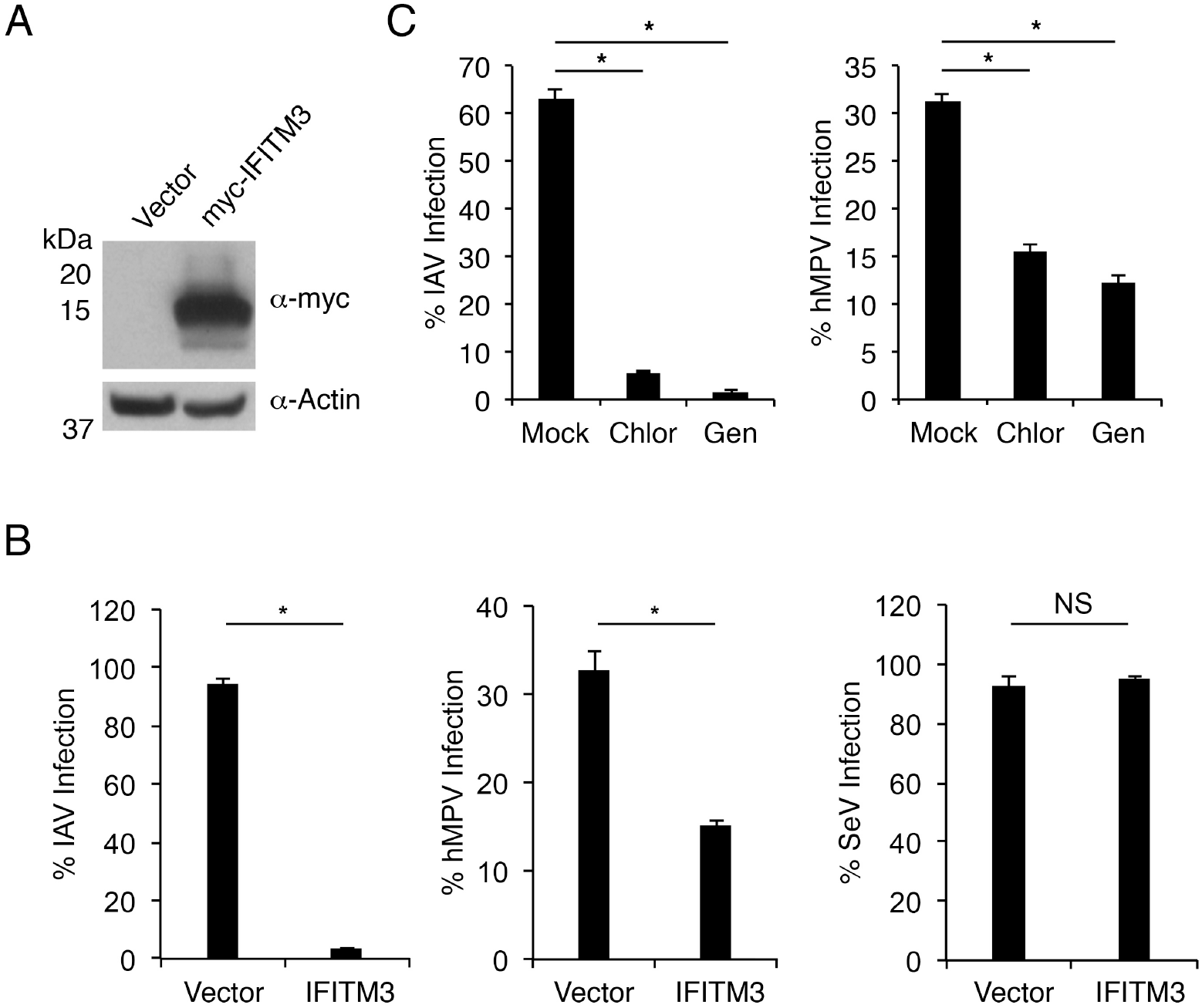
Endocytosis inhibitors and overexpressed IFITM3 restrict hMPV infection. **A)** Anti-myc and anti-Actin Western blotting of HEK293T cells stably transduced with vector control or myc-IFITM3. **B)** HEK293T cells stably transduced with vector control or myc-IFITM3 (IFITM3) were infected with IAV (MOI 2.5), hMPV (MOI 2), or SeV (MOI 2) for 24 h. Percent infection was determined by flow cytometry after staining with virus-specific antibodies using non-infected cells to set gates. Results shown are averages of three or more experiments with error bars representing standard deviation. **C)** HEK293T cells were treated for 1 h with chlorpromazine (Chlor, 10 ug/mL) or genistein (Gen, 200 uM), or were mock treated, followed by infection with IAV (MOI 2.5) or hMPV (MOI 2) in the presence or absence of the inhibitors as indicated for 24 h. Percent infection was determined by flow cytometry after staining with virus-specific antibodies using non-infected cells to set gates. Results shown are representative of three experiments with IAV, two experiments with WT hMPV and two experiments with hMPV-GFP. Error bars represent standard deviation of triplicate samples. **B,C)** Asterisks indicate p<0.001 and NS indicates not significant by Student’s t-test.

### Endogenous cellular IFITM3 limits hMPV infection

We next examined the role of endogenously expressed IFITM3 in hMPV infection using IFITM3 KO HAP1 cells (Figure 2A). Infection of IFITM3 KO cells with hMPV was significantly increased with and without IFN-α or -β treatment as compared to WT cells (Figure 2A,B, Supplementary Figure 2). Likewise, human A549 lung epithelial cells, exhibited a significant increase in susceptibility to hMPV when IFITM3 was knocked down using siRNAs in both mock and IFN-treated cells (Figure 2C,D). These results demonstrate that endogenous IFITM3 restricts hMPV infection and that other IFN-induced effectors cannot compensate for loss of IFITM3.

**Figure 2:**
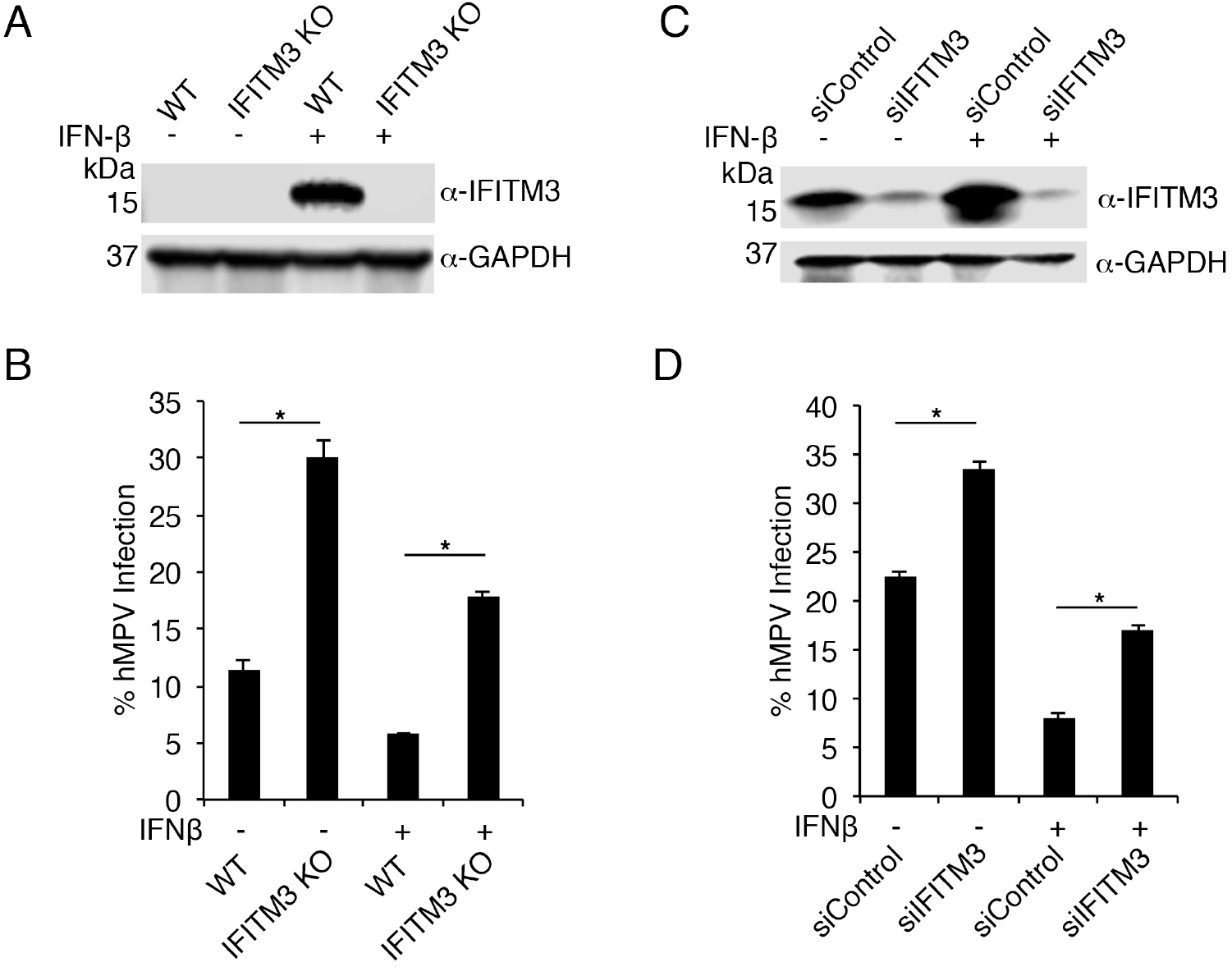
Endogenous IFITM3 restricts hMPV infection. **A,B)** WT and IFITM3 KO HAP1 cells were mock treated or treated with IFN-β overnight. **A)** Anti-IFITM3 and anti-GAPDH Western blotting. **B)** Cells were infected with hMPV (MOI 2) for 24 h. Percent infection was determined by flow cytometry after staining with virus-specific antibodies using non-infected cells to set gates. Results shown are averages of three or more experiments with error bars representing standard deviation. **C,D)** A549 cells were mock treated or treated with IFN-β overnight. Cells were simultaneously transfected with siRNA targeting IFITM3 (siIFITM3) or nontargeting control siRNA (siControl) as indicated. **C)** Anti-IFITM3 and anti-GAPDH Western blotting. **D)** Cells were infected with hMPV for 24 h. Percent infection was determined by flow cytometry after staining with virus-specific antibodies using non-infected cells to set gates. Results shown are averages of triplicate samples from an experiment representative of two experiments with WT hMPV and one experiment with hMPV-GFP. Error bars represent standard deviation. **B,D)** Asterisks indicate p<0.001 by Student’s t-test.

### Amphotericin B reverses protective effects of IFITM3

Amphotericin B is a membrane destabilizing antifungal drug that negates the protective effects of IFITM3 against IAV[22]. We sought to confirm these results and to determine whether amphotericin B could similarly reverse IFITM3 restriction of hMPV. We pretreated IFITM3-expressing HEK293T cells or vector control cells with amphotericin B, and subsequently measured infection with IAV or hMPV. We observed potent restriction of IAV by IFITM3 that was completely ablated by amphotericin B (Figure 3A). Likewise, hMPV restriction by IFITM3 was also completely reversed when cells were treated with amphotericin B (Figure 3B). These results may suggest that IAV and hMPV are inhibited by IFITM3 via similar mechanisms, and also that amphotericin B may be clinically detrimental for hMPV-infected patients as has been previously suggested for IAV infections[22].

**Figure 3:**
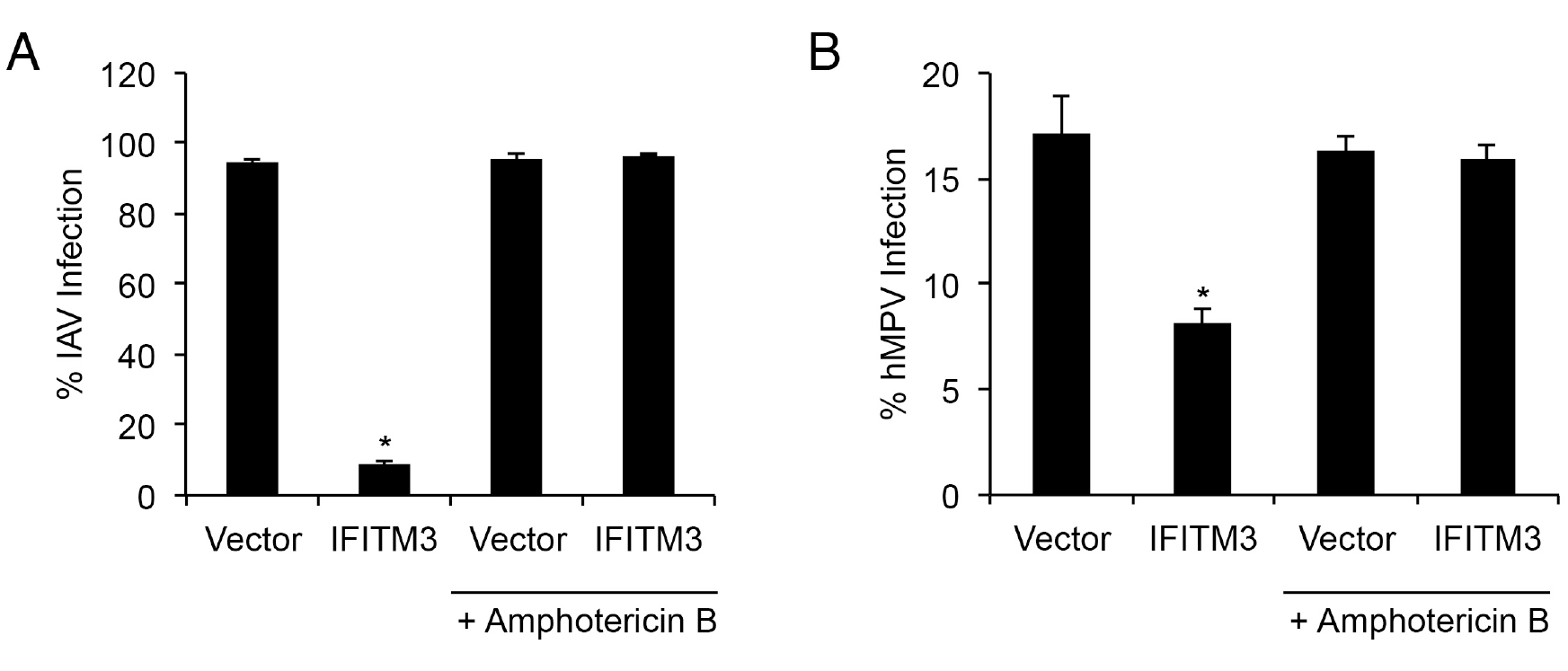
Amphotericin B reverses IFITM3 inhibition of hMPV infection. **A,B)** HEK293T cells stably transduced with vector control or myc-IFITM3 (IFITM3) were treated for 1 h with amphotericin B at a concentration of 2.5 ug/mL or were mock treated, followed by infection with IAV **(A)** or hMPV **(B)** in the presence or absence of amphotericin B as indicated for 24 h. Percent infection was determined by flow cytometry after staining with virus-specific antibodies using non-infected cells to set gates. Results shown are averages of three experiments with error bars representing standard deviation. Asterisks indicate p<0.001 as compared individually to all other samples on the respective graphs by Student’s t-test.

### Mutation of the IFITM3 endocytic trafficking motif enhances hMPV restriction

IFITM3 possesses a four amino acid YxxΦ endocytosis signal at residues 20-23 (20-YEML-23) that mediates its trafficking to endosomes and lysosomes from the plasma membrane[19, 20, 27]. A polymorphism in the human IFITM3 gene that is linked to severe influenza virus infections has been proposed to disrupt this motif by altering RNA splicing[28]. Multiple laboratories have demonstrated that mutation of Y20 to Ala within this motif results in accumulation of IFITM3 at the plasma membrane and diminishes its ability to inhibit IAV when expressed at low levels[19, 20, 27]. When strongly expressed, IFITM3-Y20A can be visualized at both the plasma membrane and at endosomes, possibly owing to passive endocytosis of the protein[20, 27, 40]. Given the dual localization of IFITM3-Y20A at the plasma membrane and at endosomes, and the proposed dual entry mechanism of hMPV, we sought to examine effects of the IFITM3-Y20A variant on hMPV infection. We thus generated a stable HEK293T cell line expressing myc-IFITM3-Y20A and confirmed that its expression was similar to cells stably expressing WT myc-IFITM3 (Figure 4A, B), and that the IFITM3-Y20A variant exhibited the expected localization compared to WT IFITM3, including plasma membrane as well as intracellular vesicle accumulation (Figure 4C). Infection of these cells with hMPV showed that IFITM3 significantly decreased infection, and remarkably, that IFITM3-Y20A expression resulted in a near-complete resistance of the cells to infection (Figure 4D). These results demonstrate that the IFITM3-Y20A variant has an enhanced ability to inhibit hMPV, correlating with its expanded pattern of localization [19, 28, 41].

**Figure 4:**
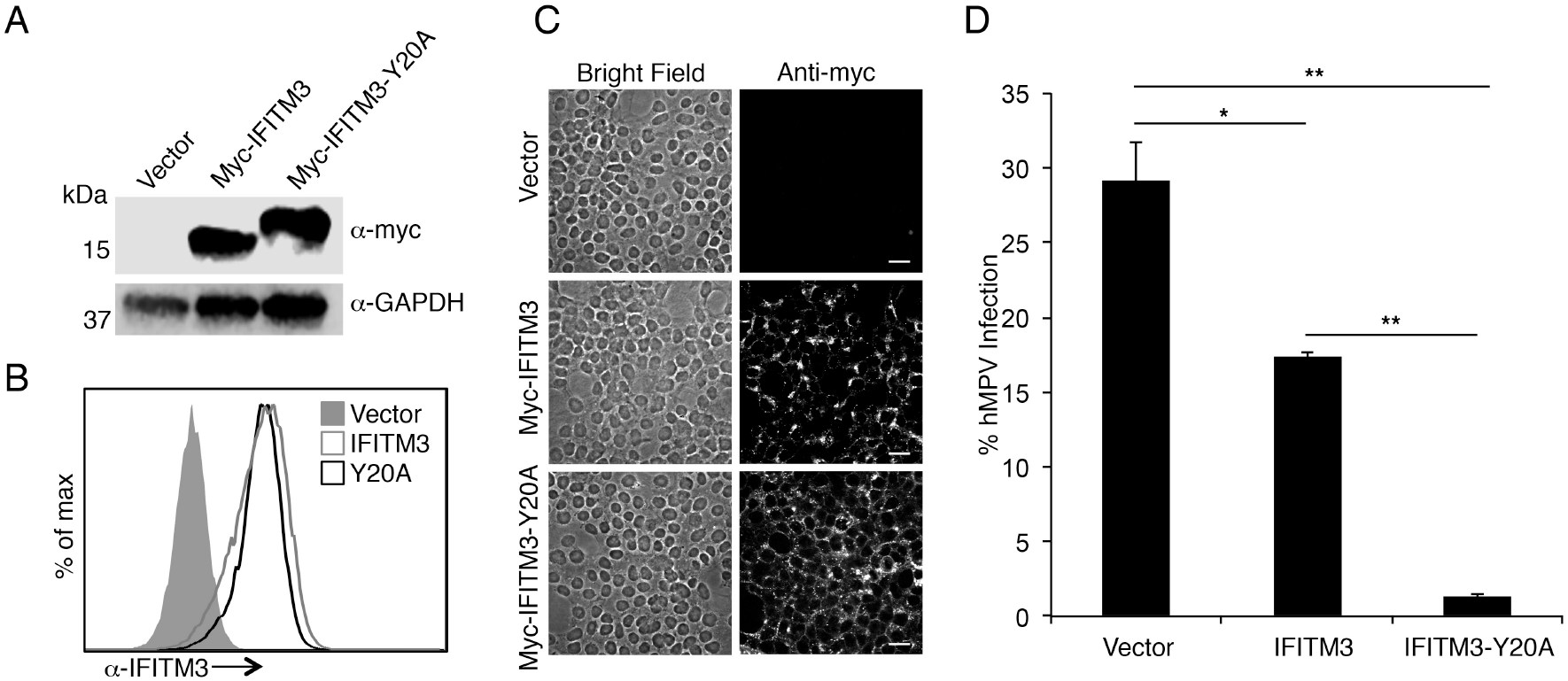
Mutation of the IFITM3 endocytosis motif enhances restriction of hMPV infection. **A)** Anti-myc and anti-GAPDH Western blotting of HEK293T cells stably transduced with vector control, myc-IFITM3, or myc-IFITM3-Y20A. **B)** Cells as in **(A)** were analyzed by flow cytometry after staining for IFITM3. C) Cells as in **(A,B)** were analyzed by bright field and confocal microscopy imaging with anti-myc staining. **D)** Cells as in **(A-C)** were infected with hMPV (MOI 2) for 24 h. Percent infection was determined by flow cytometry after staining with virus-specific antibodies using non-infected cells to set gates. Results shown are averages of four experiments with error bars representing standard deviation. Asterisk indicates p<0.001 and double asterisks represent p<0.0001 by Student’s t-test.

### Expression of IFITM3 restricts cell-cell fusion mediated by the hMPV F protein

IFITM3 prevents formation of fusion pores between IAV and host membranes[16, 17]. Thus, we reasoned that restriction of hMPV by IFITM3 likely also involves inhibition of fusion. Previous work by other groups has demonstrated that the hMPV fusion glycoprotein (F protein) is sufficient for virus membrane fusion and that it promotes cellcell fusion when expressed alone in cells[11, 12]. Previous studies have also utilized cellcell fusion assays to examine the ability of IFITMs to block fusion mediated by viral fusogens[18, 24]. Here, we used a characterized cell-cell fusion assay in which cells expressing the hMPV F protein were mixed with target cells expressing or lacking IFITM3[24]. Each population of cells was also transfected with distinct plasmids that produce luciferase only when they are present in the same cell, i.e., only when cell-cell fusion has occurred, such that fusion can be quantified by measurement of luciferase activity (Figure 5A). We first confirmed that hMPV F is capable of inducing fusion between HEK293T cell populations (Figure 5B). Importantly, fusion was only observed when hMPV F was present. Previous work by others has demonstrated that low pH is not required, but can increase fusion mediated by some hMPV F proteins[11, 12]. Indeed, pulsing of our mixed cell population with media at pH 5.0 resulted in enhanced luciferase production indicative of increased cell-cell fusion (Figure 5B). To examine effects of IFITM3 on hMPV F-mediated fusion, we transfected IFITM3, IFITM3-Y20A, or vector control into target cells. Consistent with past reports in which IFITM3 inhibited cell-cell fusion mediated by numerous viral fusion proteins[18, 24], WT IFITM3 inhibited F-mediated fusion both with a pH 5.0 pulse (Figure 5B) and at neutral pH (Supplementary Figure 4). The Y20A mutant, consistent with its increased accumulation at the plasma membrane (Figure 4C) [19, 20, 27, 41], further decreased fusion both with a pH 5.0 pulse (Figure 5B) and at neutral pH (Supplementary Figure 4). These data demonstrate that IFITM3 is able to block membrane fusion mediated by the hMPV F protein.

**Figure 5:**
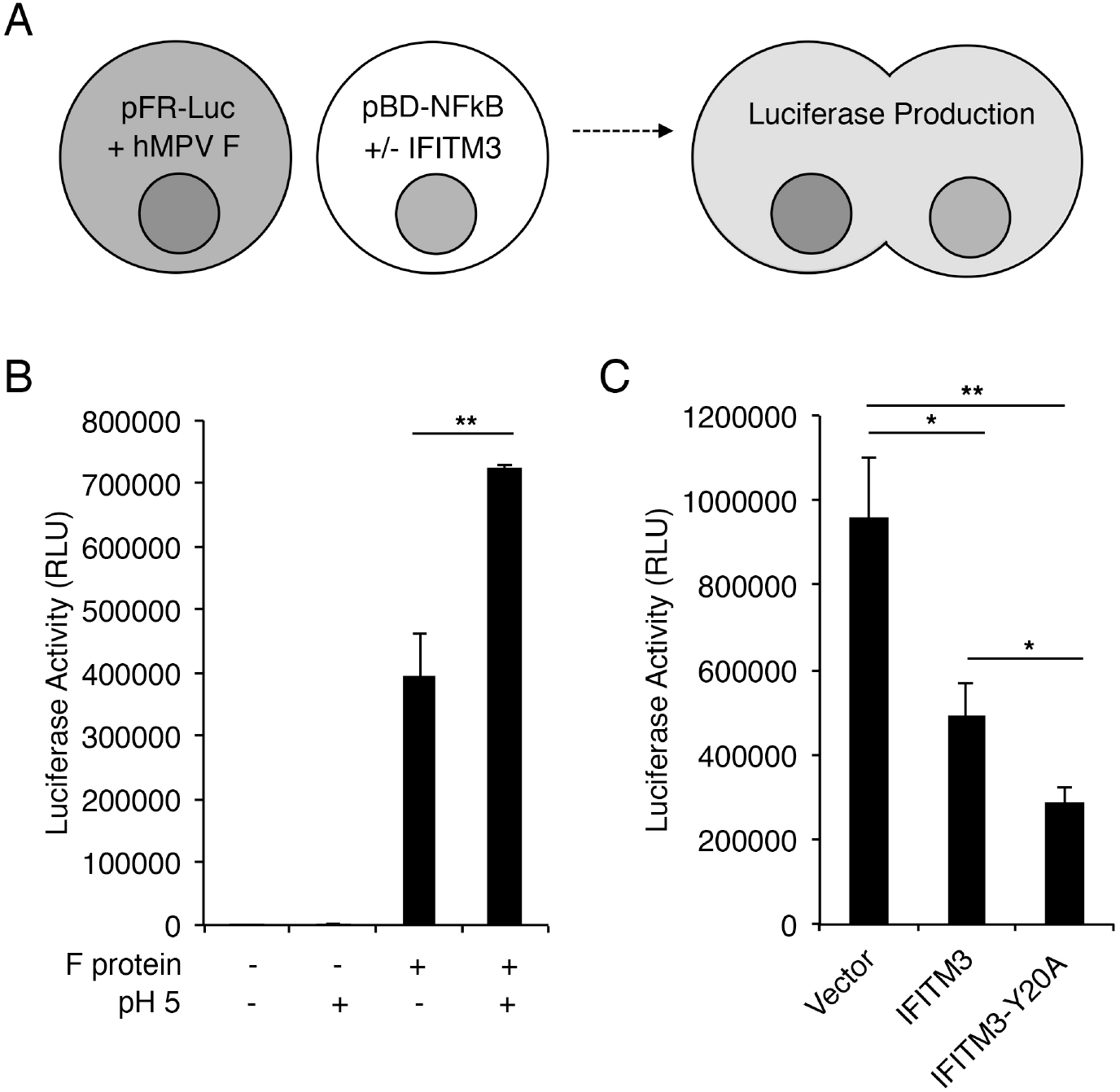
IFITM3 can inhibit membrane fusion mediated by the hMPV F Protein. **A)** Scheme for cell-cell fusion assay. **B)** HEK293T cells were transfected with pFR-Luc plus plasmid expressing hMPV F protein. A second set of cells were transfected with pBD-NFκB. Mixed cell populations were treated with media at pH 5 or with standard media for 2 min. After 8 h, cell lysates were analyzed for luciferase activity indicative of cell-cell fusion. Bar graphs depict averages of triplicate samples representative of three experiments with error bars representing standard deviation. **C)** HEK293T cells were transfected with pFR-Luc plus plasmid expressing hMPV F protein. A second set of cells were transfected with pBD-NFκB plus plasmid expressing myc-IFITM3 (IFITM3), myc-IFITM3-Y20A (IFITM3-Y20A) or vector control. Mixed cell populations were treated with media at pH 5 for 2 min. After 8 h, cell lysates were analyzed for luciferase activity indicative of cell-cell fusion. Bar graphs depict averages of triplicate samples representative of three or more experiments with error bars representing standard deviation. **B,C)** Asterisks indicate p<0.001 and double asterisks indicate p<0.0001 by Student’s t-test.

### Knockout of NEDD4 reduces hMPV infection

We previously identified NEDD4 as the primary E3 ubiquitin ligase responsible for ubiquitination and turnover of steady-state IFITM3[32]. Depletion of NEDD4 results in reduced ubiquitination of IFITM3 and increased IFITM3 levels even in the absence of infection or IFN stimulation. As such, NEDD4 KO cells are more resistant to IAV infection dependent on their enhanced IFITM3 levels[32]. Using WT or NEDD4 KO mouse embryonic fibroblasts, we confirmed increased levels of basal IFITM3 in the KO versus WT cells (Figure 6A). Upon infection with hMPV, we observed that NEDD4 KO cells were significantly more resistant to hMPV infection than WT cells (Figure 6B). These results further support a role for IFITM3 as a cellular restriction factor able to inhibit hMPV infection and also identify NEDD4 as a novel target for inhibiting hMPV infections.

**Figure 6:**
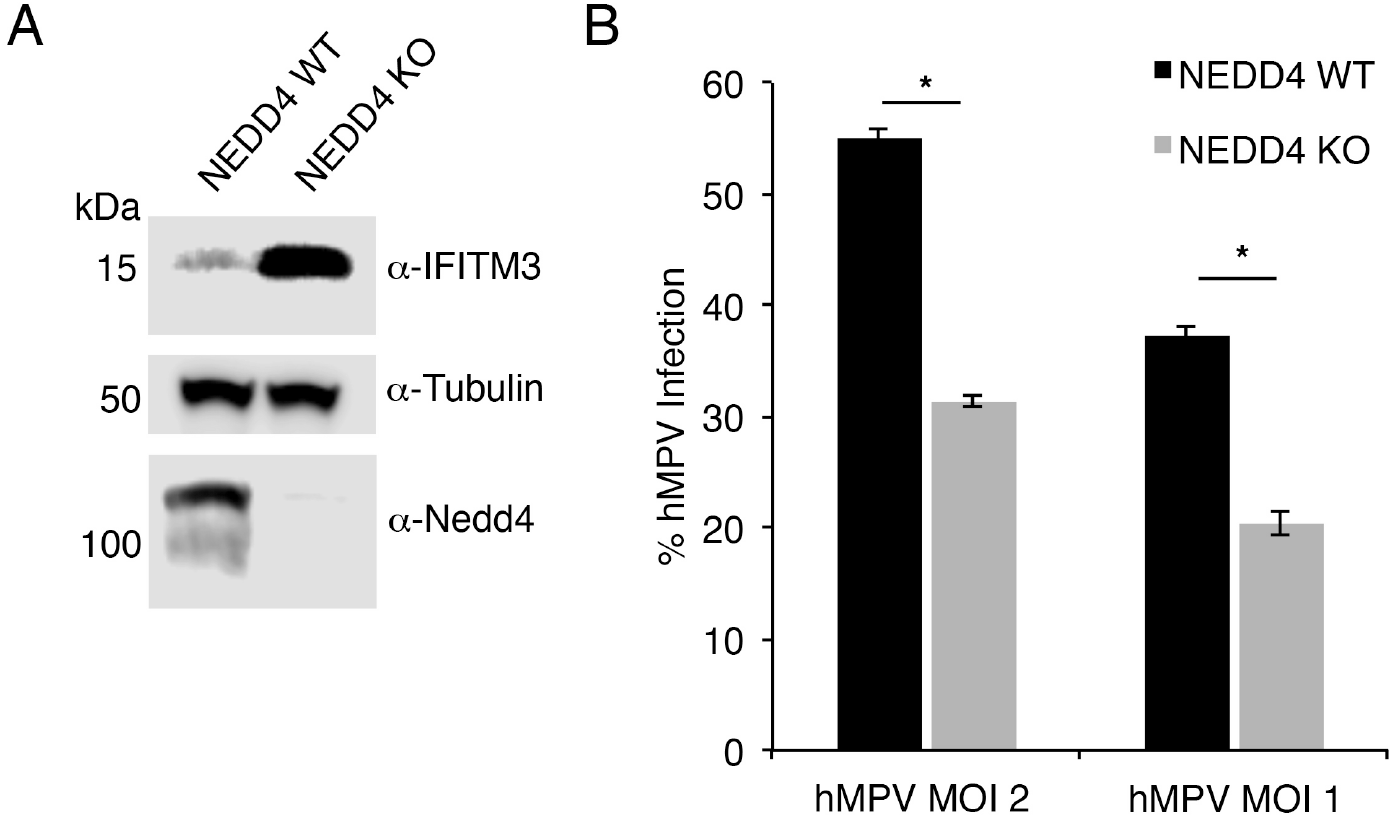
NEDD4 KO cells are partially resistant to hMPV infection. **A)** Anti-IFITM3, anti-Tubulin, and anti-NEDD4 Western blotting for WT or NEDD4 KO MEFs. **B)** Cells as in **(A)** were infected with hMPV at the indicated MOIs for 24 h. Percent infection was determined by flow cytometry after staining with virus-specific antibodies using non-infected cells to set gates. Results shown are an average of triplicate samples from an experiment representative of two experiments with WT hMPV and one experiment with hMPV-GFP. Error bars represent standard deviation. Asterisks indicate p<0.0001 by Student’s t-test.

### Toll-like receptor 7 promotes hMPV infection

In a previously published screen of more than 300 potential antiviral restriction factors, overexpression of Toll-like receptor (TLR) 7, an endosome-localized TLR that is usually associated with an antiviral response, was surprisingly observed to increase infection with an hMPV A2 strain to 131% when normalized to infection of control cells [42]. Given that IFITM3 and TLR7 are both endosomal proteins that may have opposing effects on hMPV infection, we sought to determine whether this unusual effect of TLR7 on hMPV was reproducible, and whether IFITM3 restricts infection in the presence of TLR7. To substantiate or refute the reported effect of TLR7 on hMPV, we first examined infection susceptibility of HEK293T cells when stably expressing TLR7 or vector control with the hMPV A1 strain used in our IFITM3 studies. Cells expressing TLR7 were infected at a significantly higher rate than control cells, independently confirming the published large-scale screen (Figure 7A). To determine whether endogenous TLR7 affects hMPV infection, we measured the infection susceptibility of WT and TLR7 KO murine macrophage cell lines. Consistent with our overexpression results, the percent infection of TLR7 KO cells was significantly decreased as compared to WT cells (Figure 7B). Infections of cells lacking TLR3, another endosomal protein, or TLR4, which localizes to the cell surface, were not significantly different from WT cells, providing additional specificity controls (Figure 7B). We next examined whether hMPV strain 02-202 (B1) was affected by IFITM3 and TLR7 expression in HEK293T cells. We found that, like the A1 strain, the B1 strain was susceptible to IFITM3 restriction (Supplementary Figure 3A). However, we observed no effect of TLR7 on the B1 strain, distinguishing it from the A1 strain used in our studies and the A2 strain used previously [42] (Supplementary Figure 3B). Finally, to determine whether IFITM3 can restrict hMPV infection in the presence of TLR7, we stably introduced either IFITM3 or vector control into HEK293T cells stably expressing TLR7 (Figure 7C). Upon infection with the hMPV A1 strain, the cells expressing TLR7/IFITM3 showed a robust decrease in the percentage of infection as compared to TLR7/Vector cells, indicating that IFITM3 maintains activity in the presence of TLR7. Overall, these results demonstrate that TLR7 facilitates cellular infection by certain strains of hMPV, though this enhancement of infection does not preclude restriction by IFITM3, thus highlighting the complex interactions between hMPV and cellular endosomes.

**Figure 7:**
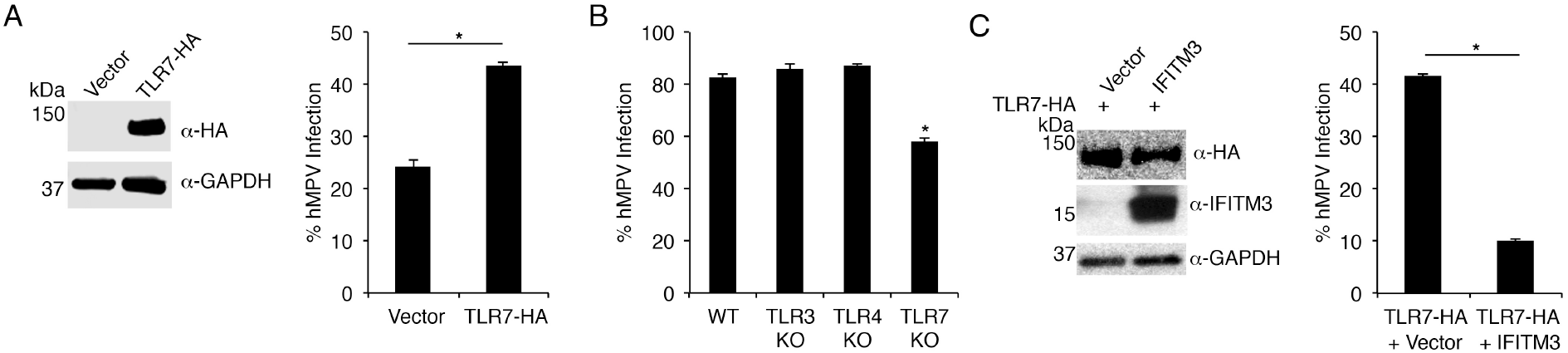
TLR7 enhances hMPV infection. **A)** HEK293T cells stably transduced with vector control or TLR7-HA were subjected to Western blotting with anti-HA and anti-GAPDH antibodies, and were infected with hMPV (MOI 2) for 24 h. Percent infection was determined by flow cytometry after staining with virus-specific antibodies using non-infected cells to set gates (bar graph). Results shown are averages of five experiments with error bars representing standard deviation of the mean. Asterisk indicates p<0.0001 by Student’s t-test. **B)** WT, TLR3 KO, TLR4 KO, or TLR7 KO murine macrophages were infected with hMPV (MOI 2) for 24 h. Percent infection was determined by flow cytometry after staining with virus-specific antibodies using non-infected cells to set gates. Results shown are averages of three experiments with error bars representing standard deviation of the mean. Asterisk indicates p<0.001 compared to all other samples in the graph by Student’s t-test. **C)** HEK293T cells stably transduced with TLR7-HA and either vector control or myc-IFITM3 were subjected to Western blotting with anti-HA, anti-IFITM3, and anti-GAPDH antibodies, and were infected with hMPV (MOI 2) for 24 h. Percent infection was determined by flow cytometry after staining with virus-specific antibodies using non-infected cells to set gates (bar graph). Results shown are averages of four experiments with error bars representing standard deviation of the mean. Asterisk indicates p<0.0001 by Student’s t-test.

## DISCUSSION

Our work has identified IFITM3 as the first confirmed restriction factor able to limit hMPV infection of cells. Infection by hMPV was decreased by overexpression of IFITM3 (Figure 1A,B) and was increased by knockout or knockdown of endogenous IFITM3 (Figure 2). Cellular pathways that regulate IFITM3 levels may thus represent new targets for limiting hMPV infections. Indeed, NEDD4 KO cells, which accumulate high levels of IFITM3 due to inefficient IFITM3 ubiquitination at steady state[32] were more resistant to hMPV infection than WT cells (Figure 6). Our data indicate that IFITM3 is able to restrict hMPV of the A1 and B1 lineages (Figure 1 and 2, and Supplementary Figure 3), and the previously mentioned large-scale overexpression screen of potential antiviral factors also showed partial inhibition of an A2 virus by IFITM3[42]. These findings indicate that IFITM3 broadly restricts multiple hMPV lineages.

Past work studying IAV and its inhibition by IFITM3 showed that the drug amphotericin B, which destabilizes lipid membranes, neutralizes the antiviral effects of IFITM3[22]. Our examination of hMPV infection of cells expressing IFITM3 showed that amphotericin B similarly negates the activity of IFITM3 against this additional virus (Figure 3B). Importantly, amphotericin B is a commonly used antifungal agent, and our new work suggests that its clinical use may enhance susceptibility to not only IAV, but also hMPV.

This work also suggests that IFITM3 inhibits hMPV infection via effects on membranes. Using cell-cell fusion assays, we found that indeed, IFITM3 is able to inhibit membrane fusion mediated by the hMPV F protein (Figure 5). This work is in line with the previously established ability of IFITM3 to block fusion pore formation in IAV infection and cell-cell fusion mediated by a multitude of viral fusion proteins[16–18, 24].

TLR7 is generally considered an antiviral molecule since it detects viral singlestranded RNA and signals for the production of type I IFNs[43]. TLR7 is expressed on macrophages, conventional dendritic cells, and plasmacytoid dendritic cells[43], and our work suggests that its presence makes these cell types more susceptible to infection with certain hMPV strains (Figure 7A,B). Staining of primary human lung airway epithelial cells with anti-TLR7 antibodies also indicated that TLR7 is expressed on these cells[44], which are additional relevant targets of hMPV infection. How hMPV utilizes TLR7 to facilitate cellular infection is not yet mechanistically understood, though the increase in the percentage of cells infected when TLR7 is expressed may suggest that TLR7 is a cell entry factor (Figure 7A,B). Remarkably, the ability of hMPV to coopt TLR7 is complemented by the reported inhibition of TLR7 signaling by the hMPV M2-2 protein[45]. Thus, some strains of hMPV have evolved to utilize TLR7 while also inhibiting its antiviral function. Detection of hMPV infection by retinoic acid inducible gene I (RIG-I), a cytosolic sensor of viral replication products, is similarly inhibited by the P or G proteins of some hMPV strains[46, 47]. Likewise, IFN signaling activity is also decreased in hMPV-infected cells via inhibition of Signal Transducer and Activator of Transcription (STAT) protein phosphorylation[48, 49]. Thus, hMPV utilizes several mechanisms to evade the type I IFN response while potentially benefitting from the presence of TLR7.

IFITM3 and TLR7 are primarily endosomal proteins, though they appear to have opposite effects on hMPV infection. In most cell types, IFITM3 levels are low prior to infection and its induction by IFNs[32]. Consistent with a role for IFN-induced proteins, such as IFITM3, in controlling hMPV infection *in vivo*, type I IFN receptor KO mice show increased hMPV titers in the lungs at early timepoints post infection[50]. It will be interesting to examine pathogenesis of hMPV infection in IFITM3 and TLR7 KO mice in the future, and to determine whether TLR7-blocking antibodies or nucleic acids can impact hMPV infections. Likewise, our work suggests that polymorphisms in the human IFITM3 gene that have been linked to severe IAV infections should also be examined for effects on hMPV infections[28–30]. Overall, we have revealed that two endosomal proteins, IFITM3 and TLR7, differentially regulate vulnerability of cells to hMPV infection, though IFITM3 appears to dominate when both proteins are expressed (Figure 7C). These findings support the proposed role for endocytosis in hMPV infection, and have identified new factors that may be manipulated or targeted to prevent or treat infection by this important human pathogen.

## Acknowledgments

We thank Dr. John Williams for providing us with the hMPV B1 strain used in this work.

## Notes

### Author Contributions

T.M.M., J.L., and J.S.Y. designed experiments and analyzed data. T.M.M., Y.Z., A.D.K., L.Z., M.L., A.Z., and M.C. performed experiments. T.M.M. and J.S.Y. wrote the manuscript with editorial contributions from all authors.

---

**Supplementary Figure 1:**
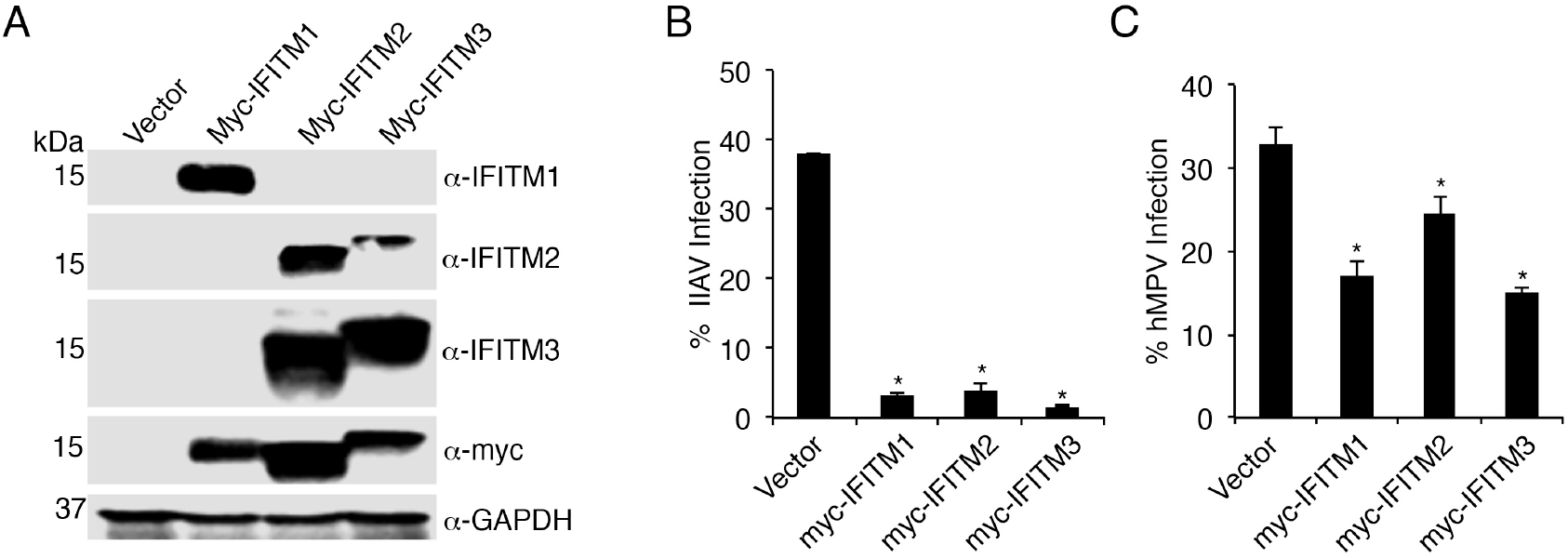
Overexpressed IFITMs restrict hMPV infection. **A)** Anti-myc and anti-GAPDH Western blotting of HEK293T cells stably transduced with vector control, myc-IFITM1, myc-IFITM2, or myc-IFITM3. **B,C)** HEK293T cells stably transduced with vector control or myc-IFITMs were infected with IAV (MOI 2.5) **(B)** or hMPV (MOI 2) **(C)** for 24 h. Percent infection was determined by flow cytometry after staining with virus-specific antibodies using non-infected cells to set gates. Results shown are averages of two or more experiments with error bars representing standard deviation. Asterisks indicate p<0.001 as compared to vector control by Student’s t-test. Some data from Main Text Figure 1 is re-plotted here for comparison between IFITM3 and other IFITMs.

**Supplementary Figure 2:**
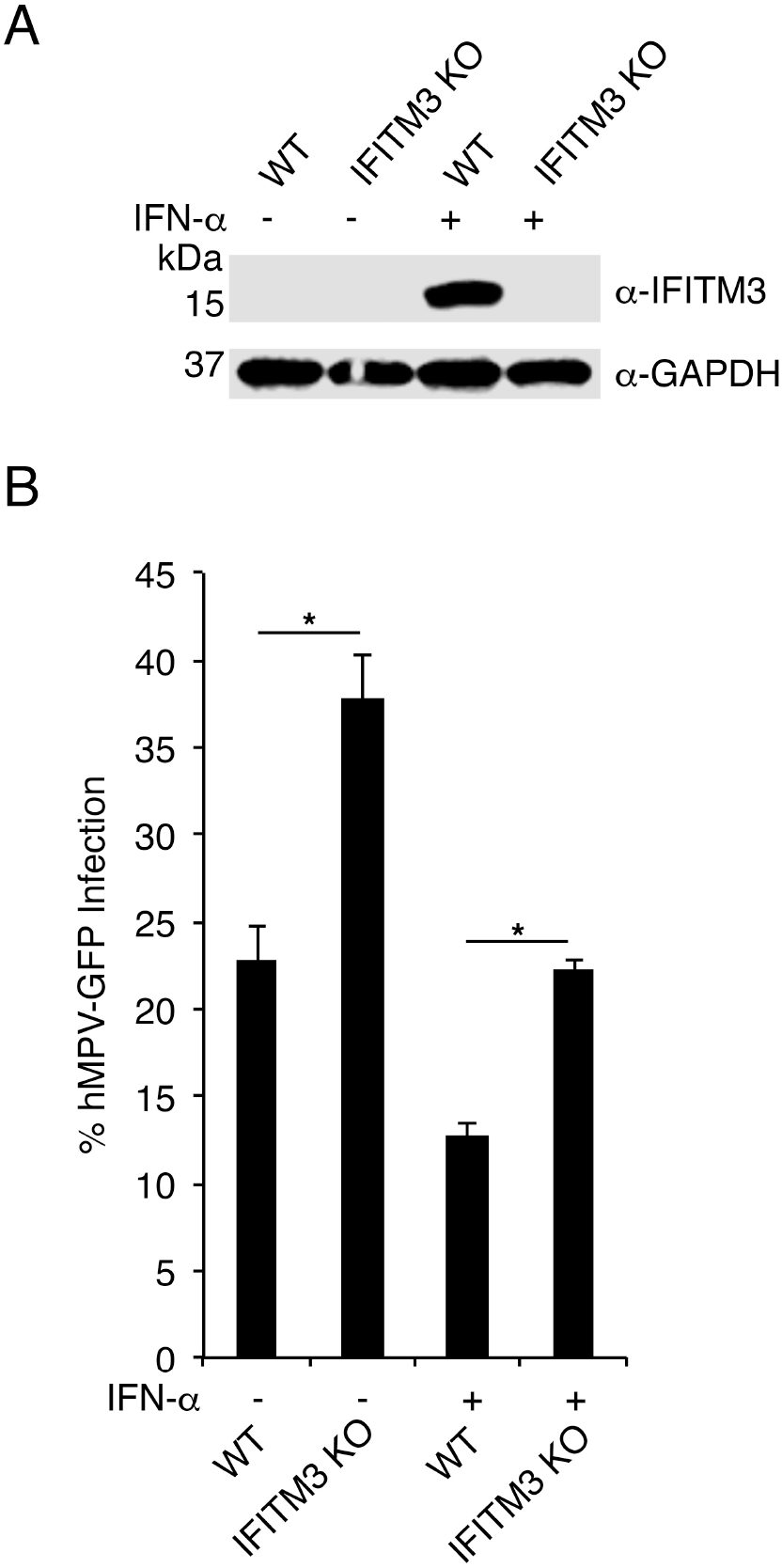
Endogenous IFITM3 restricts hMPV infection. **A,B)** WT and IFITM3 KO HAP1 cells were mock treated or treated with IFN-α overnight. **A)** Anti-IFITM3 and anti-GAPDH Western blotting. **B)** Cells were infected with hMPV-GFP (MOI 2) for 24 h. Percent infection was determined by flow cytometry detection of GFP using non-infected cells to set gates. Results shown are representative of two experiments with error bars representing standard deviation of triplicate samples. Asterisks indicate p<0.001 by Student’s t-test.

**Supplementary Figure 3:**
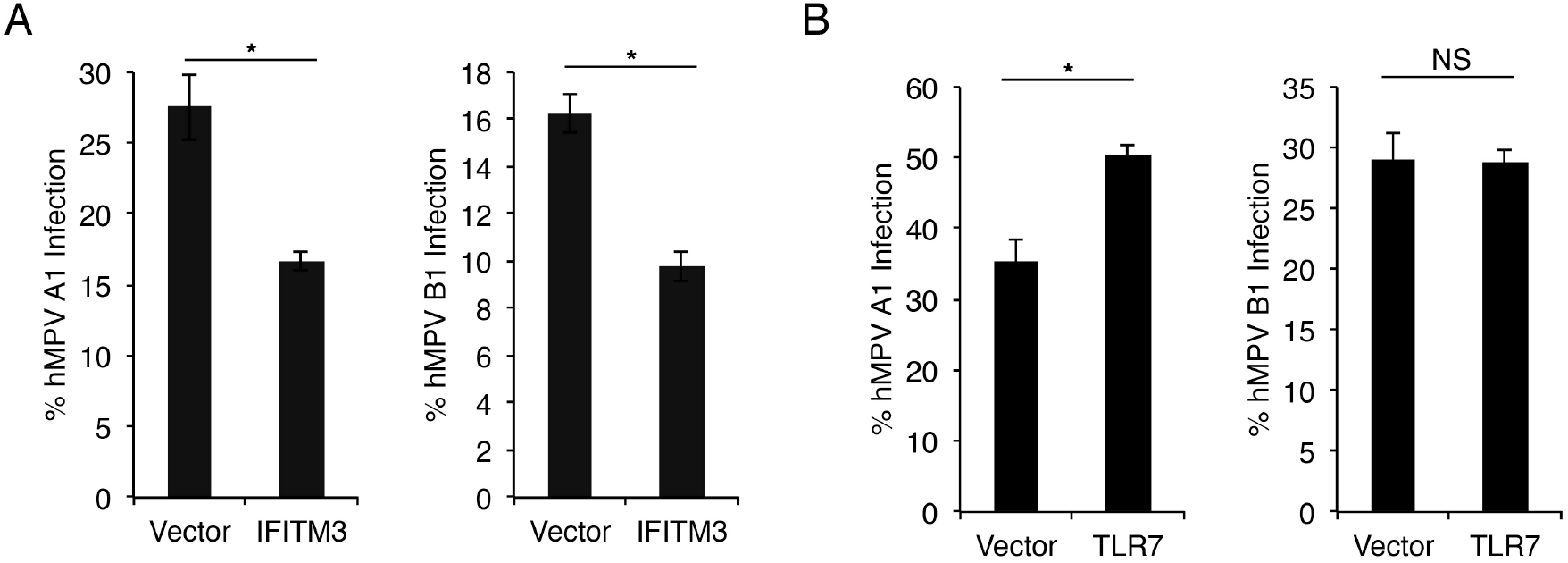
Strain-specific effects of IFITM3 and TLR7 on hMPV infection. **A)** HEK293T cells stably transduced with vector control or myc-IFITM3 (IFITM3) were infected with hMPV A1 or B1 strains (MOI 2) for 24 h. Percent infection was determined by flow cytometry after staining with virus-specific antibodies using non-infected cells to set gates. Results shown are averages of triplicate samples from an experiment representative of more than three experiments with error bars representing standard deviation. **B)** HEK293T cells stably transduced with vector control TLR7-HA (TLR7) were infected with hMPV A1 or B1 strains for 24 h. Percent infection was determined by flow cytometry after staining with virus-specific antibodies using non-infected cells to set gates. Results shown are averages of triplicate samples from an experiment representative of more than three experiments with error bars representing standard deviation. **A,B)** Asterisks represent p<0.001 and NS indicates not significant by Student’s t-test.

**Supplementary Figure 4:**
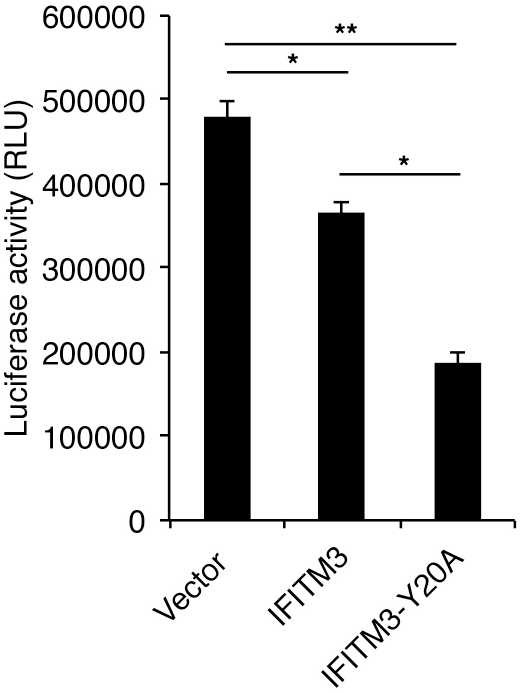
IFITM3 can inhibit membrane fusion mediated by the hMPV F Protein. HEK293T cells were transfected with pFR-Luc plus plasmid expressing hMPV F protein. A second set of cells were transfected with pBD-NFκB plus plasmid expressing myc-IFITM3 (IFITM3), myc-IFITM3-Y20A (IFITM3-Y20A) or vector control. The two cell populations were mixed and plated for 8 h and cell lysates were analyzed for luciferase activity indicative of cell-cell fusion. Bar graphs depict averages of triplicate samples representative of two experiments with error bars representing standard deviation. Asterisks indicate p<0.001 and double asterisks indicate p<0.0001 by Student’s t-test.

